# RNA-specific condensation promotes auto-regulation of polyQ RNA-binding protein Whi3

**DOI:** 10.1101/2021.04.25.441362

**Authors:** Therese M. Gerbich, Erin M. Langdon, Grace A. McLaughlin, Benjamin M. Stormo, Blair Yu, Marcus Roper, Amy S. Gladfelter

**Author notes:** These authors contributed equally on this work (Erin M. Langdon, Grace A. McLaughlin).

## Abstract

RNA-binding proteins are frequently seen to be capable of auto-regulation by binding their own transcripts. In this work, we show that in the multinucleated fungus *Ashbya gossypii*, the phase-separating RNA-binding protein Whi3 binds and regulates its own transcripts in distinct condensates from its other targets. Failure of Whi3 to bind its own transcript leads to a reduction in Whi3 protein level and its inability to properly regulate its other targets, leading to defects in nuclear cycling, polarized growth, and transcription of START-regulated genes. These results present a role for auto-regulation and condensate formation triggered by an RNA-binding protein interacting with its own coding transcript. Given the propensity of RNA-binding proteins to interact with their own coding mRNAs, this may be a wide-spread mechanism of feedback that utilizes biomolecular condensates.

## Introduction

RNA-binding proteins (RBPs) frequently are able to bind to their own mRNA (Müller-McNicoll et al. 2019). In some cases, this allows them to function as part of autoregulatory feedback loops to control their own expression which in turn can impact other cellular processes in which they participate (Müller-McNicoll et al. 2019). Broadly speaking, there are two ways by which such autoregulatory feedback loops can function. In the first case, an RBP binding to its own mRNA will repress the further production of that RBP. This creates a negative feedback loop wherein high levels of a particular RBP lead to a homeostatic control mechanism that will prevent further accumulation of that RBP (Müller-McNicoll et al. 2019). In mammalian cells, for example, SF2/ASF negatively regulates its own expression by binding to its own transcript to promote alternative splicing of that transcript into a version that does not code for the full-length isoform (Sun et al. 2010). A second mode of action for autoregulatory feedback loops between RBPs and their mRNA transcripts is through positive feedback that amplifies an input signal into a robust switch-like response (Müller-McNicoll et al. 2019). An example of this is seen with the *Drosophila* protein Orb, which is required in flies for establishment of the dorsoventral and anteroposterior axes (Tan et al. 2001). Orb promotes the localization and translation of its own mRNA first at the dorsal anterior part of the oocyte, and then at the posterior pole. The autoregulatory positive feedback loop ensures that there will be sufficient quantities of Orb protein at the correct time and place to act on its other mRNA targets that are required to establish these axes during development (Tan et al 2001).

RNA-binding proteins are increasingly linked to biomolecular condensates in which RNA promotes the demixing of the proteins and it is likely that these auto-regulatory processes could be regulated through condensation (Roden & Gladfelter, 2021). The RNA-binding protein Whi3 forms condensates to regulate multiple processes in the syncytial fungus *Ashbya gossypii*, including polarized growth and control of the nuclear cycle (Lee et al. 2013; Lee et al. 2015). Whi3 condenses with its target mRNAs from these processes to exert local control over the nuclear cycle and polarized growth (Lee et al. 2013; Lee et al. 2015). Specifically, Whi3 forms condensates with the cyclin mRNA *CLN3* near nuclei to regulate nuclear cycling and Whi3 protein forms distinct condensates with the formin mRNA *BNI1* near hyphal tips to regulate polarized growth (Lee et al. 2013; Lee et al. 2015). Condensation is sensitive to levels of both Whi3 protein and its mRNA targets (Zhang et al. 2015), and specificity is achieved through mRNA secondary structure elements that allow certain mRNA to incorporate into particular Whi3 droplets. For example, in addition to *BNI1, SPA2* (another Whi3 target mRNA whose protein product is important for normal polarized growth) transcripts are able to partition into polarity droplets, but *CLN3* mRNA is excluded (Langdon et al. 2018).

While Whi3 was originally discovered for its role in regulating *CLN3* in *S. cerevisae* (Nash et al. 2001), it is now known to bind a large number of mRNA targets (Cai et al. 2013). Though a comprehensive list of all Whi3 targets in *Ashbya* has not been published, we reasoned that it likely has more targets that have yet to be discovered. Indeed, there are a number of mRNAs in the genome of Ashbya that have an enrichment of the consensus sequence binding site for Whi3; most notably the *WHI3* mRNA. While we have a growing understanding on how Whi3 regulates nuclear cycling and polarized growth in *Ashbya* distinctly at a local level, we hypothesized that Whi3 might also regulate its own transcript to create feedback loops to strengthen local control of the different cellular processes it regulates. We reasoned that perhaps Whi3 protein could be condensing with *WHI3* mRNA to create or repress production of Whi3 protein to tune local levels of Whi3 in order to regulate other local Whi3 targets, similar to what has been seen with Orb. In this study, we find that Whi3 does interact with *WHI3* transcript, can condense with its own RNA and that this interaction is required for Whi3 to perform its normal roles in regulating *CLN3* and *BNI1*. These experiments provide a link between biomolecular condensation and auto-regulation of RNA-binding proteins, a major constituent of condensates in cells.

## Results and Discussion

We observed that *WHI3* mRNA contains five instances of UGCAU (Figure 1A), the consensus binding site for Whi3 protein, and a similar density of binding sites as known Whi3 targets such as *CLN3* (Riorden et al. 2011). Given this, we hypothesized that Whi3 was binding and regulating its own transcript to promote cell polarity and asynchronous nuclear division in *Ashbya*. In order to test this idea, we created a *whi3-5m* allele (which we call *wbsm*) where all the Whi3 binding sites were removed from *WHI3* transcript via silent mutations (Figure 1A). To assess if Whi3 interaction with these sites is relevant to the LLPS capacity of Whi3 protein, we *in vitro* transcribed wild-type *WHI3* and *wbsm* RNA and combined the synthesized RNAs with recombinantly-expressed Whi3 protein. Notably, Whi3 protein was able to phase separate with *WHI3* mRNA but not with *wbsm* mRNA (Figure 1B), suggesting Whi3 is capable of using these predicted binding sites to bind and phase separate with its own mRNA. The protein demixes in a protein and RNA concentration dependent manner, similar to other known RNA targets of Whi3 (Figure 1C). These data support that these sequences are relevant sites of interaction and are required to drive Whi3 protein to undergo LLPS in vitro.

**Figure 1.**
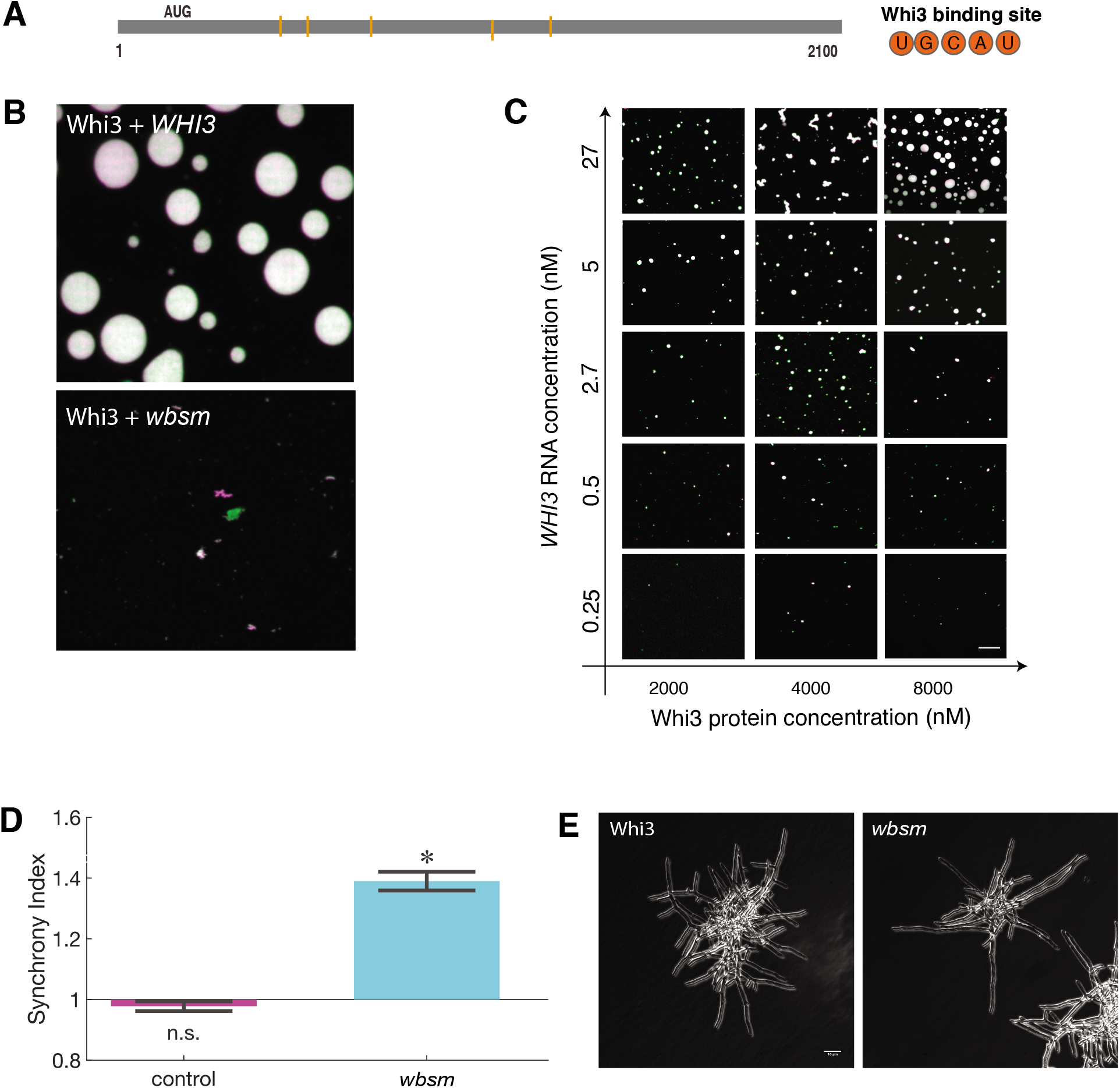
Whi3 binds and phase separates with its own transcript. (A) *WHI3* contains 5 UGCAU predicted Whi3 binding sites. (B) Representative images phase separation assays of Whi3 with either *WHI3* or *wbsm* mRNA. Whi3 protein concentration 8000 nM, RNA concentration 5 nM. (C) Phase diagram of Whi3 and *WHI3*. (D)) Synchrony indices for *wbsm* as compared to control cells. n>175 for all strains. Bars denote SE. (E) Representative images of cell morphology in control and *wbsm* cells. Scale bar 10 µm.

To assess the functional relevance of these sequence elements, we next replaced *WHI3* in *Ashbya* cells at the endogenous locus with the *wbsm* construct and tagged the mutant protein with td-Tomato. We first looked to see if the *wbsm* cells had altered cell cycle or polarity phenotypes. We found that these cells were more synchronous in their nuclear division cycles than control cells as assessed by scoring the cell cycle state of neighboring nuclei and comparing to what would be expected to be found by chance (Figure 1D) (Gerbich et al. 2020). The *wbsm* cells had a synchrony index around 1.4, indicating a moderate phenotype that is less severe than what is seen in the *whi3ΔpolyQ* or the *whi3Δ*, which are around 1.7 and 1.9, respectively (Nair et al. 2010; Lee et al. 2015). Additionally, the *wbsm* mutants had defects initiating new lateral branches during polarized growth (Figure 1E). These phenotypes were similar to *whi3* mutants that cannot bind mRNA (*whi3ΔRRM*), cannot form Whi3 assemblies (*whi3ΔpolyQ*), or complete Whi3 null mutants (*whi3Δ*) (Lee et al. 2015). Importantly, the *wbsm* mutant encodes a completely wild-type protein indicating that changes in the RNA-sequence alone were sufficient to block function, supporting the idea that Whi3 binding its own transcript is functionally important.

In order to understand how Whi3 binding its own transcript is impacting Whi3 function, we then looked to see if there was a change in Whi3 protein localization in *wbsm* cells. We found that compared to control cells, *wbsm* cells had a reduced number of Whi3 puncta around nuclei and at tips (Figure 2A-C), consistent with the defective cell cycle and polarity phenotypes observed.

**Figure 2.**
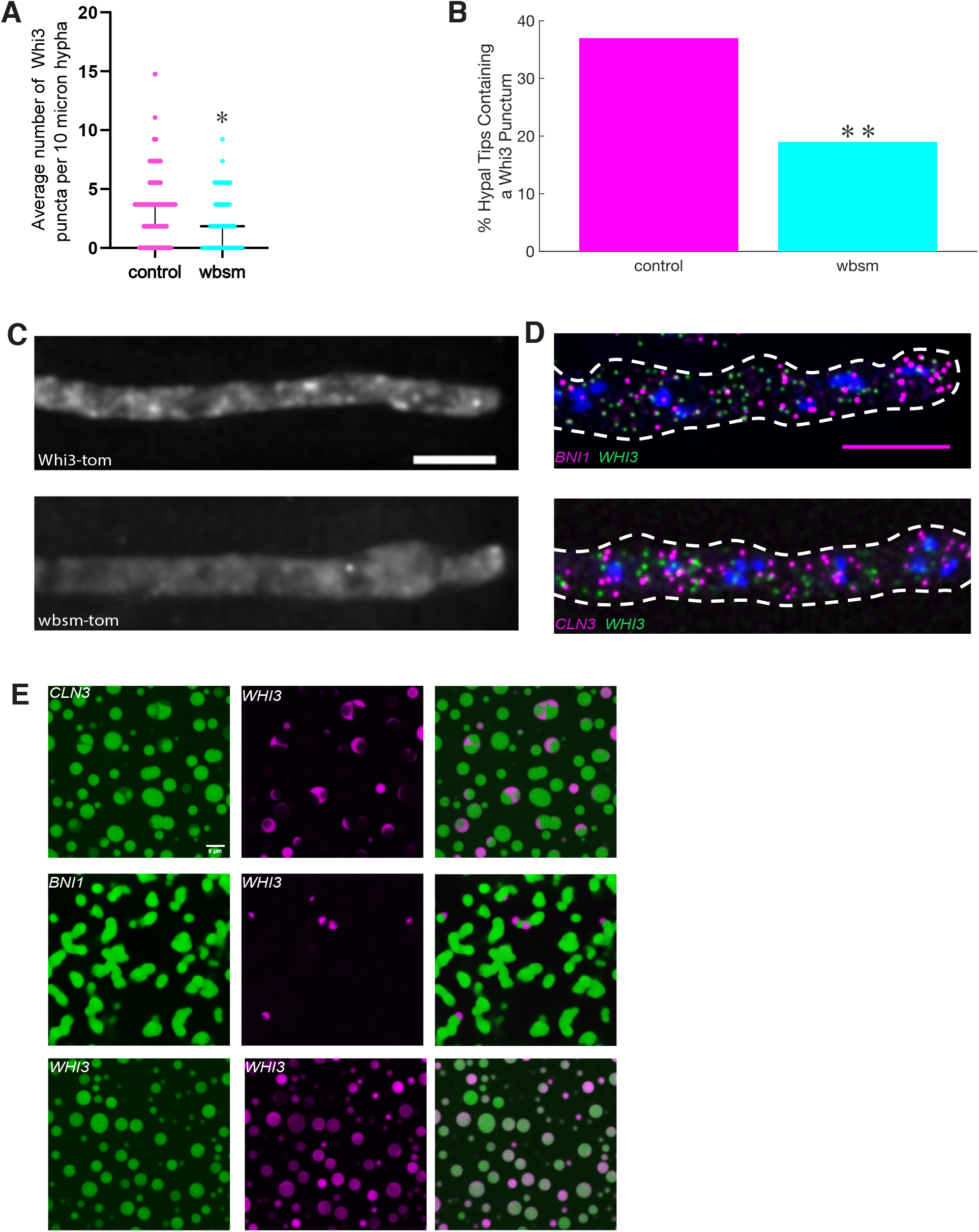
*WHI3* condensates are distinct from *CLN3* or *BNI1* condensates. (A) Average number of Whi3 puncta per 10 µm hyphal segment in control and wbsm strains. \**p* < 0.05 by t-test. Bars 95% CI. n>70 for each strain. (B) Percent of hyphal tips in control and wbsm strains containing a Whi3 punctum. \**p* < 0.05 by N-1 Chi-Square test. n>70 for each strain. (C) Representative images of Whi3-tom in *Ashyba* control or wbsm cells. Images are max projections. Scale bar 5 µm. (D) Representative images of *WHI3* and *BNI1* or *WHI3* and *CLN3* in *Ashyba* control cells. Images are max projections. Scale bar 5 µm. (E) Representative images of phase separation assays where *WHI3* RNA was added to pre-formed droplets of Whi3 protein that were initiated with either *CLN3, BNI1*, or *WHI3* (labeled in each panel). Scale bar 5 µm.

Given the loss of function phenotypes and reduced numbers of Whi3 puncta around nuclei and at tips, we predicted that *WHI3* mRNA is a necessary component of normal *CLN3* and *BNI1* droplets in the cells. However, when we used sm RNA F.I.S.H. to look at *WHI3* transcripts alongside *CLN3* or *BNI1*, we found that the mRNAs did not colocalize suggesting they are in distinct cellular condensate compartments (Figure 2D). Consistent with the lack of colocalization in cells, *WHI3* mRNA did not co-localize with either *CLN3* or *BNI1* droplets in vitro. Synthesized *WHI3* mRNA was not able to partition into Whi3 protein droplets containing *CLN3* or *BNI1* mRNAs (and vice versa) *in vitro*, but was able to partition into pre-formed droplets composed of Whi3 protein and *WHI3* mRNA (Figure 2E). These data indicate that Whi3 protein’s ability to associate with its own RNA is required for normal formation and function of Whi3 condensates in cells but this role is not the result of *WHI3* RNA being a core component of *CLN3* and *BNI1* droplets but rather functions as a separate condensate.

Given that *WHI3* does not appear to be a component of *CLN3* or *BNI1* droplets under normal circumstances in wild type cells, we reasoned that perhaps the cell cycle and polarity phenotypes we observed were due to changes in Whi3 protein levels. Whi3 protein abundance was indeed impacted in our *wbsm* cells. We found that by fluorescence microscopy (Fig 3A) and Western blot (Fig 3B), Whi3 protein levels were reduced in these cells by about 50%.

**Figure 3.**
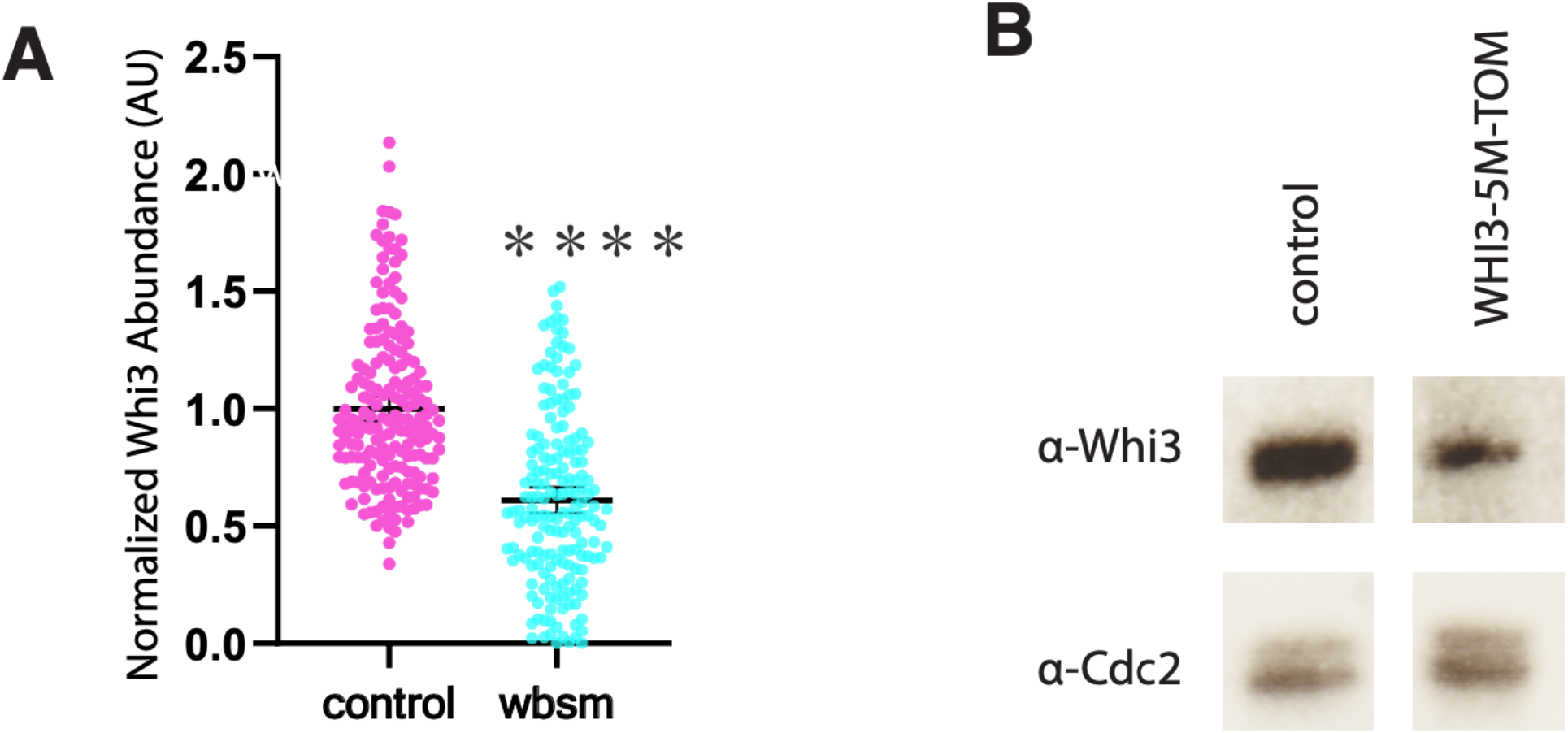
Whi3 protein levels are reduced in *wbsm* cells. (A) Normalized Whi3 abundance in control and *wbsm* mutant cells. \**p* < 0.05 by t-test. Bars 95% CI. n>175 for all strains. (B) Western blot of Whi3 abundance in control and wbsm cells. Cdc2 is a loading control.

We suspect therefore that the cell cycle and polarity phenotypes that we see in these cells are due to reduced availability of Whi3 protein to form condensates with and to regulate *CLN3* and *BNI1*.

How would silent mutations in *WHI3* lead to lowered abundance of Whi3 protein? We had noticed that upstream of the *WHI3* ORF, there were four predicted MBP binding sites (ACGCGN) (Figure 4A). In the closely related budding yeast *S. cerevisiae*, passage through Start is regulated by MBF protein binding to these sites. This occurs downstream of Cln3 in the genetic circuit, suggesting to us that perhaps in *Ashbya* Whi3 bound to its own mRNA to regulate levels of Whi3 protein and control passage through Start as part of an autoregulatory feedback loop. In this model, when Cln3 promotes passage through start, it also promotes the transcription of *WHI3*, which leads to increased local levels of Whi3 protein, which in turn leads to even more *CLN3* and thus Cln3, ensuring commitment to passage through start via a positive feedback loop. Consistent with this, in *wbsm* cells there are fewer *CLN1/2* transcriptional hotspots (Figure 4C), suggesting that in these cells passage through start is impaired. Importantly, we also found that in *wbsm* cells, there were fewer nuclei transcribing *WHI3* which suggests that less of it is being produced (Figure 4 B&C).

**Figure 4.**
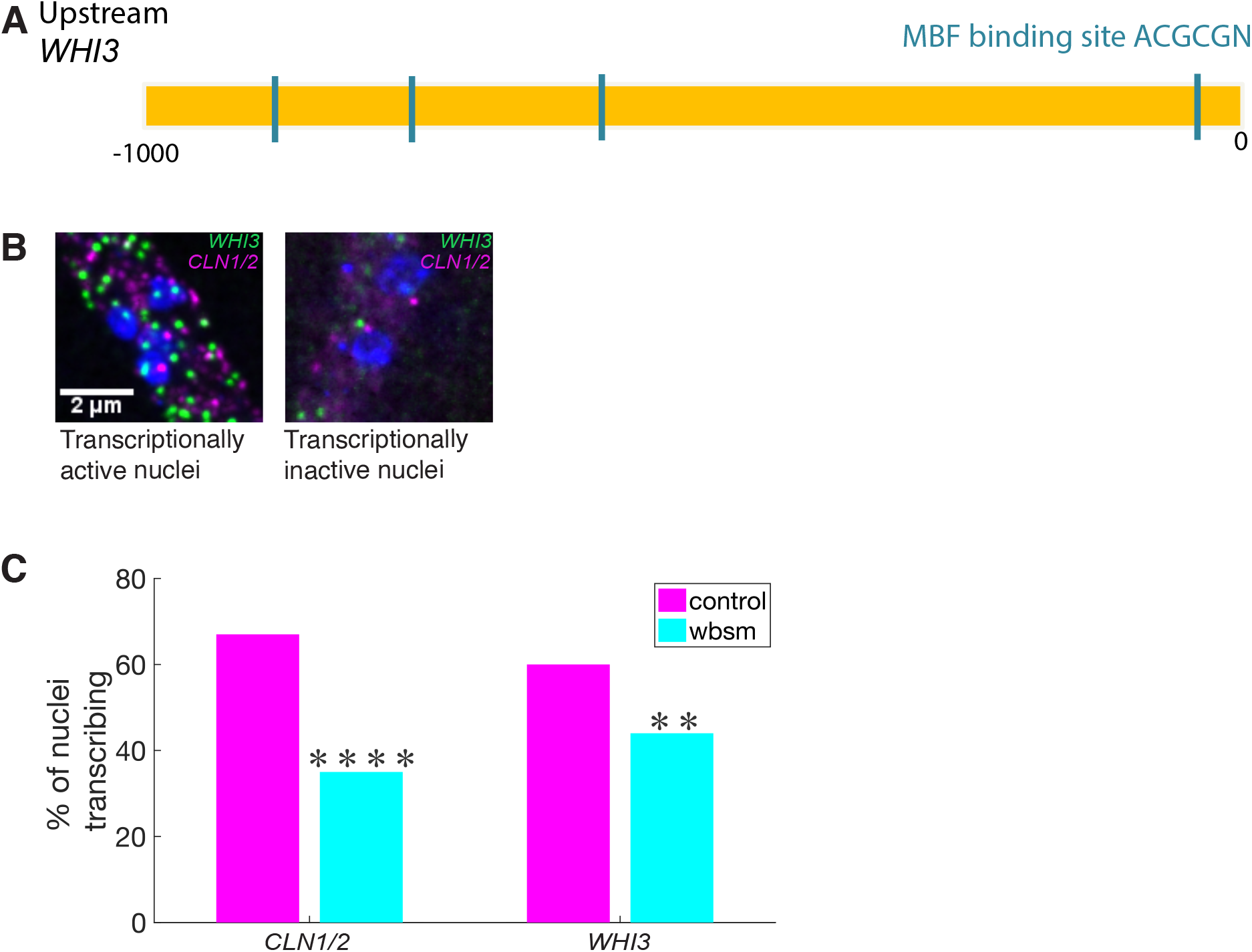
*CLN1/2* and *WHI3* transcription are reduced in *wbsm*. (A) Upstream of *WHI3* are 5 ACGCGN predicted MBF binding sites. (B) Representative images of transcriptionally active and inactive nuclei. Scale bar 2 µm. (C) Percent of nuclei transcribing *CLN1/2* or *WHI3* in control or *wbsm* cells. \**p* < 0.05 by N-1 Chi-Square test. n>140 for all strains.

## Conclusion

We found that Whi3 binds and regulates its own transcript in *Ashbya gosspii* cells. When Whi3 cannot bind its own transcript, Whi3 protein levels are decreased and the cells are no longer able to undergo normal nuclear cycling and polarized growth. Furthermore, in these cells transcription of Start-regulated genes is decreased, suggesting that perhaps Whi3 autoregulation is important for controlling passage through Start in *Ashbya* via an autoregulatory positive feedback loop. We predict that Whi3 autoregulates its own transcripts by stabilizing the RNAs and/or promoting their local translation via condensation.

## Acknowledgements

We would like to thank the Gladfelter Lab for useful discussions. The work was supported by National Institutes of Health grant R01-GM-081506, the HHMI faculty scholars program, and NSF grant MCB-1840273. The authors declare no competing financial interests.

## Materials and Methods

### Strain Construction

To create *wbsm* mutant strain tagged with tomato, a geneblock containing the mutations of interest were ordered from IDT (Table 1). This geneblock was then used to replace the relevant portions of Whi3 in AGB993 using Gibson cloning to amplify the geneblocks and relevant portions of the plasmid (Table 2). Constructs were sequenced across the Whi3 region of the plasmid to ensure that no additional mutations had accrued during the cloning process. Each plasmid was then cut by SacII, XhoI, and NdeI. The 6.377 kb fragments containing the mutant Whi3, the fluorescent tag, and the NAT resistance marker were separated and purified using an agarose gel and integrated into *Ashbya* using protocols described previously (Wendland et al., 2009).

**Table 1.**
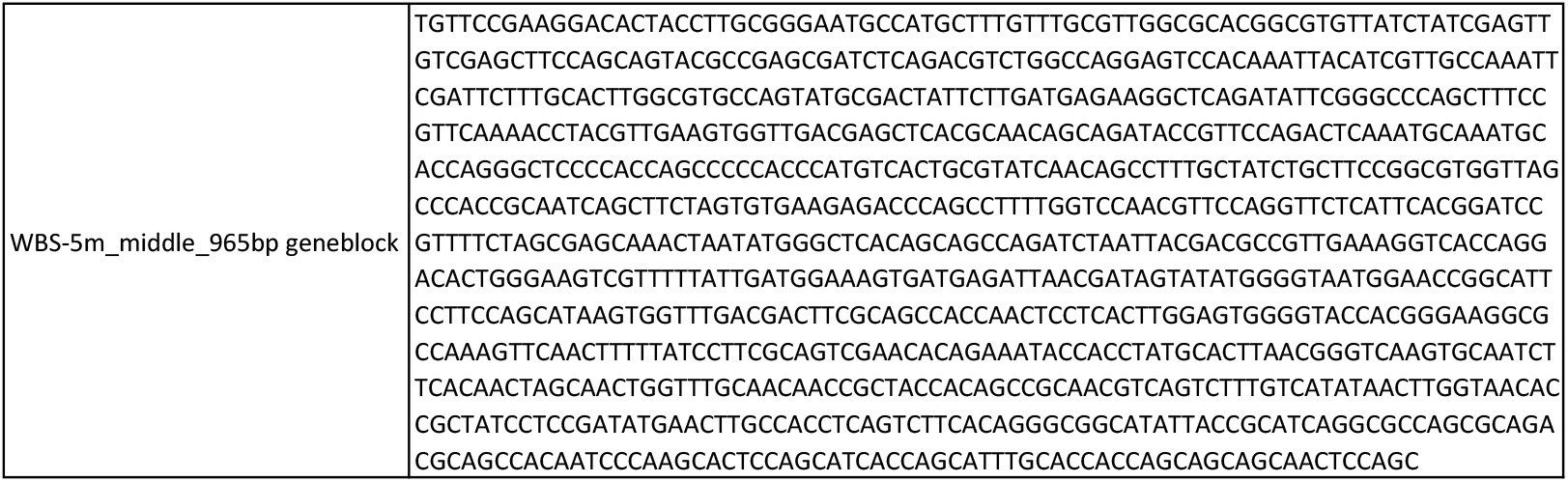
Geneblock used to create *wbsm*.

**Table 2.**
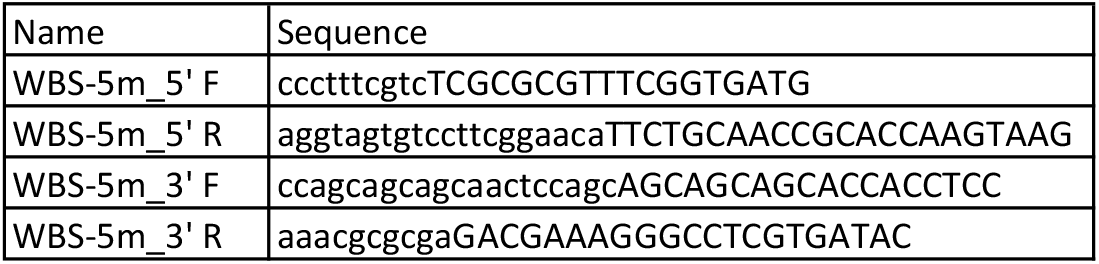
Primers used to create *wbsm*.

### Recombinant protein purification

Full length Whi3 with an N-terminal 6-His tag was transformed into BL21 bacterial cells for protein expression. Overnight cultures were used to inoculate 1L cultures, and protein expression was induced when cultures reached OD 0.6 using 1 mM isopropyl β-d-1-thiogalactopyranoside. Cultures were grown while shaking for 20h at 18°C. Cells were collected via centrifugation and lysed in lysis buffer (1.5M KCl, 20 mM Tris pH 8.0, 20 mM Imidazole pH 8, 1 mM DTT, 1 tablet of Roche protease inhibitor cocktail). Sonication was performed to further lyse cells and to shear DNA. Lysis was then clarified by further centrifugation and supernatant was incubated with Ni-NTA resin. Beads were washed with 20 column volumes of lysis buffer in a gravity column before being eluted in elution buffer (150 mM KCl, 20 mM Tris pH 8.0, 200 mM Imidazole pH 8.0,1 mM DTT). Immediately following elution, proteins were dialzed into droplet buffer (150 mM KCl, 20 mM Tris pH 8.0, 1 mM DTT) overnight at 4°C and then used to perform phase separation assays.

### In vitro RNA transcription

RNA transcription was perfomed as described previously (Langdon et al 2018). Briefly, plasmid DNA was digested to obtain a linear DNA template that was used with a T7 Hiscribe in vitro transcription kit and cy3-UTP or cy5-UTP according to manufacturer’s instructions to create labeled versions of *CLN3, BNI1, WHI3*, and *wbsm*.

### In vitro phase separation assays

Phase separation assays for Whi3 and *WHI3* or *wbsm* were performed by diluting RNA and protein into droplet buffer to the desired final concentration in glass chambers. Reactions were incubated for 2h at RT and imaged using a spinning disc confocal microscope (Nikon CSU-W1) with VC Plan Apo 60X/1.40 NA oil immersion objective and an sCMOS 85% QE camera (Photometrics).

For mixing experiments, initial droplets were pre-formed according to protocol above. After 2h, 5mM *WHI3* was added and allowed to mix for 30 min. Imagining was performed as described in above.

### Synchrony Imaging and Index

*Ashbya* cells were grown under appropriate selection in Ashbya full media (AFM) for 16h shaking at 30°C. 3.7% formaldehyde was added for fixation and cells were returned to 30°C for 1 hr. Following fixation, cells were washed thoroughly with PBS and resuspended in Solution A (100 mM K2PHO4 pH 7.5, 1.2M Sorbitol). Zymolysase digestion at 30°C was used to remove cell wall, with samples being checked visually every 10 minutes to monitor digestion progress. After digestion, cells were washed thoroughly in Solution A and then spotted onto polylysine treated wells. Cells were washed with PBS and blocked with BSA, and then incubated with rat α-alpha tubulin antibody overnight at 4°C. Cells were then washed with PBS and incubated with fluorescently labeled secondary antibody and Hoechst. After a final washing with PBS, cells were covered with Prolong gold mounting medium and a coverslip was sealed onto the slide for imaging. Imaging was performed using a widefield microscope (Nikon Eclipse TI stage) with a Plan Apo λ 100x/1.45 Oil Ph3 DM objective and an Andor Zyla VSC-06258 camera. Images were deconvolved using 25 iterations of the Lucy-Richardson algorithm in Nikon Elements software and then processed using Fiji. Cell synchrony index was performed as described previously (Nair et al., 2010). Briefly, it is a method developed by our lab in collaboration with statisticians allowing us to compare the likelihood that adjacent nuclei are in the same cell cycle state due to local synchrony compared to what would be expected by chance.

### Cell Branching Imaging

*Ashbya* cells were grown for 14h shaking in *Ashbya* full media (AFM) at 30°C with appropriate selection. Cells were imaged using a widefield microscope (Nikon Eclipse TI stage) with a Plan Apo λ 60x/1.40 Oil Ph3 DM objective and an Andor Zyla 4.2 plus VSC-06258 camera.

### Whi3 protein imaging and Analysis

*Ashbya* cells were grown under appropriate selection in AFM for 12.5h shaking at 30°C. Cells were collected by centrifugation and washed with 2X low fluorescence minimal media (LFM). Cells were resuspended in LFM and transferred to a 1.4% agarose gel pad made with LFM on a slide. Coverslips were sealed with VALAP and cells were immediately imaged using a widefield microscope (Nikon Eclipse TI stage) with a Plan Apo λ 60x/1.40 Oil Ph3 DM objective and an Andor Zyla 4.2 plus VSC-06258 camera. Images were deconvolved using 12 iterations of the Lucy-Richardson algorithm in Nikon Elements software and then processed for display using Fiji.

To measure Whi3 abundance by fluorescence, hyphal segments were outlined in the phase channel in a middle plane to create ROIs. The ROIs were then used on Z-sum projections of the fluorescent channel to determine the average fluorescent signal in each hyphal ROI. These values were then averaged to determine the Whi3 abundance in each strain. wbsm protein level was compared to control cells using a T-test.

To measure hyphal tips containing a Whi3 punctum, hyphal tips were marked in the phase channel, followed by manual scoring of the fluorescent channel for the presence or absence of a Whi3 punctum. wbsm was compared to control cells using an N-1 Chi-Square test. To determine the average number of hyphal Whi3 puncta, hyphal segments of 10 µm were outlined in the phase channel, followed by manual scoring of the number of Whi3 puncta in that region in the fluorescent channel. wbsm was compared to control cells using a T-test.

### Western Blot

*Ashbya* cells were grown under appropriate selection in Ashbya full media (AFM) for 16h shaking at 30°C. Cells were spun down and rinsed with PBS before being resuspended in 2× NP-40 buffer (6 mM Na2HPO4, 4 mM NaH2PO4 × H2O, 1% NP-40, 150 mM NaCl, 2 mM EDTA, 50 mM NaF, 4 µg/ml leupeptin, 0.1 mM Na3VO4 + fresh 1× EDTA-free protease inhibitor, 1 mM PMSF, 0.01 mg/ml leupeptin, and 50 mM β-glycerophosphate). Cells were flash frozen with liquid nitrogen followed by lysis using a commercial coffee grinder at 4°C. Lysate was clarified by centrifuging for 10 minutes at 13.K rpm and 4°C. Lysate concentration was normalized using NP-40 buffer, and lysate was run in a 10% polyacrylamide gel followed by western blotting. Anti-Whi3 is a custom polyclonal antibody raised in rabbits and affinity purified, and was used at 1:1000. Anti-cdc2 is the commercial monoclonal antibody Anti-PSTAIR and was also used at 1:1000.

### smFISH Imaging

RNA smFISH labeling of Ashbya was performed as previously described (Lee et al., 2013). Briefly, *A. gossypii* cells were grown under appropriate selection in AFM for 16h shaking at 30°C. 3.7% formaldehyde was added for fixation and cells were returned to 30°C for 1 hr. Following fixation, cells were collected by centrifugation and washed with cold Solution A before being resuspended in spheroplasting buffer (100 mM K2PHO4 pH 7.5, 1.2M Sorbitol, 2mM Vanadyl ribonucleoside complex). Zymolyase digestion to remove cell wall was performed as above, at 30°C with samples being checked visually every 10 minutes to monitor digestion progress.

Subsequent steps were perfomed in RNAse free conditions. Following digestion, cells were washed with cold Solution A and incubated in 70% EtOH at 4°C for between 6 and 16h. Cells were then washed gently with SSC wash buffer and resuspended in hybridization buffer (100 mg/mL dextran sulfate, 1µg/mL *E. coli* tRNA, 2mM Vanadyl ribonucleoside complex, 0.2 mg/mL BSA, 2X SSC, 10% v/v deionized formamide). TAMRA or cy5 conjugated RNA FISH probes (Stellaris LGC Biosearch Technologies) complementary to RNA of interest were added and cells were incubated overnight at 37°C. Cells were then gently washed again with SSC wash buffer and incubated with Hoechst for 30 min at room temperature. Cells were gently washed for a final time with SSC wash buffer and then mounted on glass slides with Prolong gold mounting medium. Coverslips were sealed to slides and slides were left to rest overnight at room temperature before imaging.

Cells were imaged using a widefield microscope (Nikon Eclipse TI stage) with a Plan Apo λ 100x/1.45 Oil Ph3 DM objective and an Andor Zyla VSC-06258 camera. For colocalization experiments, Images were deconvolved using 29 iterations of the Lucy-Richardson algorithm in Nikon Elements.

### Analysis of Transcriptionally Active Nuclei

To measure transcriptionally active nuclei, max projections of images were created and manually scored for the presence of mRNA hotspots (Dundon et al. 2016). *wbsm* strain was compared to control using an N-1 Chi-Square test.

